# A flexible framework for minimal biomarker signature discovery from clinical omics studies without library size normalisation

**DOI:** 10.1101/2024.07.03.601811

**Authors:** Daniel Rawlinson, Chenxi Zhou, Kim-Anh Lê Cao, Lachlan J.M. Coin

## Abstract

Application of transcriptomics, proteomics and metabolomics technologies to clinical cohorts has uncovered a variety of signatures for predicting disease. Many of these signatures require the full ‘omics data for evaluation on unseen samples, either explicitly or implicitly through library size normalisation. Translation to low-cost point-of-care tests requires development of signatures which measure as few analytes as possible without relying on direct measurement of library size.

To achieve this, we have developed a feature selection method (Forward Selection-Partial Least Squares) which generates minimal disease signatures from high-dimensional omics datasets with applicability to continuous, binary or multi-class outcomes. Through extensive benchmarking, we show that FS-PLS has comparable performance to commonly used signature discovery methods while delivering signatures which are an order of magnitude smaller. We show that FS-PLS can be used to select features predictive of library size, and that these features can be used to normalize unseen samples, meaning that the features in the complete model can be measured in isolation for making new predictions.

By enabling discovery of small, high-performance signatures, FS-PLS addresses an important impediment for the further development of precision medical care.

## Introduction

The goal of providing accurate disease diagnosis from high-throughput ‘omics assays is being actively pursued by biologists, computational scientists, and diagnostics companies. The omics assays in question - transcriptomics or proteomics for instance - are rich in information and typically capture many thousands of features to describe the biological state of the patient or donor. Indeed, the number of features measured usually exceeds the number of samples collected in a study by several fold, which presents difficulties for optimal fitting of coefficients to the features included in the model (Xu & Jackson, 2019).

A machine learning (ML) approach to diagnostics development seeks to take the high-dimensional data generated with an omics assay and generate a mapping from the input feature space to an output probability or prediction of disease status. As computing power has accelerated and complex ML models have become more accessible, much of the application of ML to biology has embraced the full feature set to train models and provide new predictions (eg. Beaude et al., 2023; Cui et al., 2024). However, feature selection, or the refinement of the feature set to a smaller number of the most relevant molecules, is still of utility in the clinical domain. Firstly, feature selection offers interpretability as to what aspects of biology are important for the predicted diagnosis (Cai et al., 2018). Secondly, it addresses the problems associated with generating a model using an overwhelming number of features against comparatively few samples (Altman & Krzywinski, 2018). And thirdly, a minimal feature set adds the possibility for conversion of the ‘omics test into a readily available manufactured test that is cheaper and faster than a complete ‘omics assay (Gliddon et al., 2018). The development of simplified testing procedures is of particular benefit for resource-constrained settings where complex laboratory procedures are impracticably onerous (Mabey et al., 2004).

Existing methods for feature selection fall into three broad categories of wrapper, filter, or embedded approaches (Saeys et al., 2007) Wrapper-based methods select individual features sequentially and fit a model after each iteration, thus allowing the algorithm to decide when a sufficient number of features have been included. Forward selection, backward selection, and recursive feature elimination all fit into this category. Filter-based methods determine a univariate measure of relevance for each candidate feature (eg. R^2^, Spearman’s ρ), and apply a threshold cut-off to include just the most relevant features into a model. Embedded approaches unite the feature selection and model building steps such that an objective function can be evaluated, and an optimal model can be analytically derived (Saeys et al., 2007). Additionally, projection-based methods have seen extensive use in the biological domain due to the very high dimensional structure of ‘omics datasets. While projection-based methods are helpful for managing high-dimensionality, these methods collapse the dimensionality down into orthogonal, latent components rather than selecting individual features for modelling, so further steps need to be taken to restrict the actual number of features (eg. Lê Cao et al., 2011).

Embedded feature selection methods using regularisation are used extensively in research on signature generation (eg. Habgood-Coote et al., 2023). Occupying this category are LASSO (Tibshirani, 1996) and Elastic-Net (Zou & Hastie, 2005), both of which solve linear models of X regressed on y with a penalty imposed on the L1 norm of the maximum likelihood coefficients. The two differ in that LASSO imposes a penalty strictly on the L1 norm, whereas Elastic-Net uses an additional mixing parameter *alpha* to modulate the proportion of the penalty meted on the L1 and L2 norms. Consequently, Elastic-Net generally produces models with more non-zero feature coefficients, but these have lower magnitude owing to the influence of the L2 portion of the penalty in an effort to limit overfitting. Many modified regularisation methods have been employed in signature discovery, such as Adaptive LASSO (Zou, 2006), Relaxed LASSO (Meinshausen, 2007), and Bayesian formulations (Park & Casella, 2008). However, the original LASSO and Elastic-Net designs remain widely used.

Minimum Redundancy Maximum Relevance (mRMR) is a filter-based method that is popular for biological signature generation because it reduces collinearity in the final feature set. mRMR ranks the candidate features by balancing two criteria: rewarding mutual information shared between the feature and the outcome variable, and penalising mutual information shared between each feature and all features preceding it (Ding & Peng, 2005). The modeller can then select the number of minimally redundant features that is desired for modelling with some other function (Li et al., 2022).

Whatever method is used for deriving a small biological signature from compositional (rather than absolute) count data, application of the signature to unseen data still requires a full ‘omics dataset for normalisation. Signatures requiring normalisation are not truly small as all features must be measured to resolve the compositional constraint of the key features in the signature. This has typically been addressed by the somewhat ad-hoc use of house-keeping genes for count normalisation, yet the optimal strategy for normalisation remains a significant impediment to the translation of biological signatures into minimal diagnostic tools.

In this work we describe Forward Selection – Partial Least Squares (FS-PLS), a novel and flexible method for feature selection from omics data that delivers small signatures for prediction of disease state in binary and multinomial classification problems. FS-PLS combines the dimensional reduction and component orthogonality of projection-based modelling methods (PLS) with the sparsity and clinical interpretability of a wrapper-based method (FS).

We demonstrate that FS-PLS generates sparse signatures with little deterioration of performance when compared with more dense models, and with improved performance in comparison with other sparse feature selection methods. Additionally, by applying FS-PLS to select features that predict library size, we show that classification models can incorporate normalisation features that do away with the need for a true library size and require only the minimal feature set for prediction of new samples. Further, the derived signatures suffer no significant loss in accuracy compared with conventionally normalised samples. In this way FS-PLS provides a unified solution to identification of complete signatures of disease, which is applicable to any compositionally-constituted ‘omics biomarker discovery problem.

FS-PLS has already been used to discover biomarkers from gene expression data (Gliddon et al., 2021; Herberg et al., 2016; Jackson et al., 2023). Here our aim is to evaluate its effectiveness against the backdrop of widely used signature generation methods. We assess FS-PLS’s performance against the embedded methods LASSO and Elastic-Net, and the filter method mRMR (minimum Redundancy-Maximum Relevance), which ranks features according to the amount of shared information with the outcome variable while penalising their shared information with other variables. We apply the methods to five distinct ‘omics datasets: two microarray, two RNA-Seq, and one proteome (see *Methods*), which are comprised of three binary and two multi-class outcome types. Additionally, we demonstrate FS-PLS’s advantage over standard forward selection approaches of Stagewise FS for binary and StepAIC for multinomial. Our comparison is focused on linear methods as these explicitly select or rank features for inclusion rather than inferring feature importance once a classifier has been made as with tree-based or neural network modelling. We follow our comparison of methods with an assessment of FS-PLS’s performance with the incorporation of normalisation features on the two RNA-Seq datasets and with a separate validation dataset for one of these.

Ultimately, our work shows that FS-PLS is a robust method for feature selection from omics data that should be considered when researchers are seeking minimal signatures of disease state. FS-PLS is available in open access at https://github.com/lachlancoin/fspls.

## Methods

### Description of FS-PLS

Denote *X* a matrix of observations (m) with variables (p) and an outcome variable *y*, which can be binary, multinomial, or continuous. The matrix *X* is centred so that the mean of each variable is 0, and the means of the variables are recorded as μ_1_, …, μ_p_. The pseudocode below describes the main steps of the algorithm:

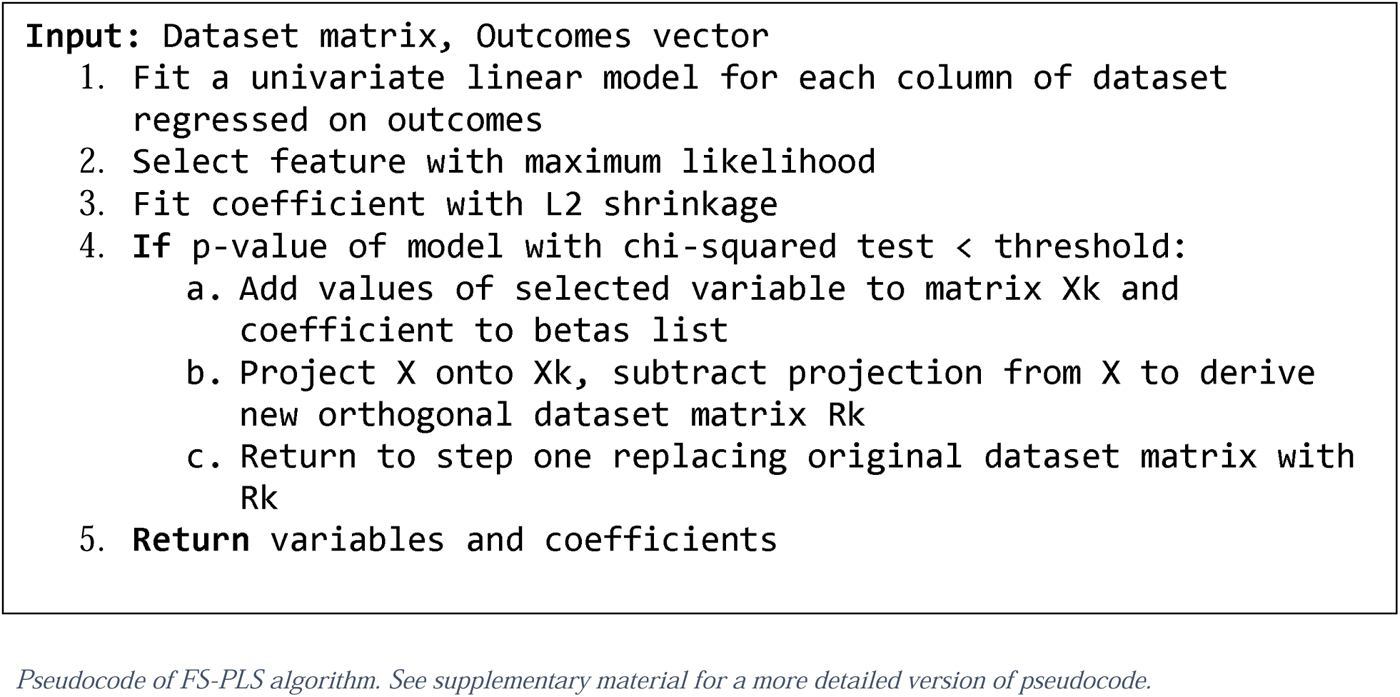

The FS-PLS algorithm proceeds in *k* iterations, where at each iteration a new variable is selected and fitted until some stopping criteria is satisfied (described below). At the *k*_th_ iteration we take *X^k^* to be the set of *k* variables already selected from the data: *X^k^* =[*x_i1_,*…,*x_ik_*], which at initialisation is a matrix of 0 columns. The Singular Value Decomposition (SVD) is applied to *X^k^* to obtain an orthonormal basis *U* on the *k*-dimensional subspace of *X*, which is spanned by the *k* selected variables of *X^k^*.

The full-dimensional *X* matrix is then projected onto *X_k_*with *P^k^* = *UU^T^X* to determine the portion of each element of *X* that is captured by the *k*-dimensional subspace. The original data *X* is subsequently deflated with *P^k^* to obtain *R^k^*= *X-P^k^*, a new *mxp* matrix whose columns are all orthogonal to the previously selected variables *X^k^*.

From here, FS-PLS fits a univariate linear model for each *j* of *R^k^* and chooses the one with the maximum log likelihood for addition into the feature set. In so doing, a new variable *j* is selected that exhibits the greatest correlation with *y* that is orthogonal to previous variables. Whereas standard forward selection might subtract the current estimate ỹ from *y* to orthogonalize the remaining variation, FS-PLS instead borrows from PLS by deflating X. This is done so that at each successive iteration, the previously selected variables are completely removed from the space of *X* and have no chance of re-entering the model at a later stage. Moreover, removing the component of X that covaries with *X^k^* means that variables that correlate with *X^k^* are also precluded from entering the model in place of a previously selected *k* at a later iteration.

Once a new variable is selected, FS-PLS re-fits its coefficient with L2 shrinkage applied as in ridge regression and the variable is added to *X^k^*for the next iteration of the algorithm. A chi-squared test is performed to determine the significance of the log-likelihood of the new variable against the null model (i.e. just the *y* vector and the offset). FS-PLS stops adding variables once the p-value exceeds a threshold, or when a pre-defined maximum number of variables has been selected.

For prediction of new data, FS-PLS maintains the coefficients for each variable in its deflated space and applies the saved orthogonalization transformations at each variable. Model estimates are thus fixed once estimated at each variable selection step, which ensures that only the portion of each variable that is uncorrelated with previous variables is incorporated into the model.

### Datasets

#### Microarray datasets

The microarray dataset from Golub *et al*. (1999) is a collection of bone marrow and peripheral blood samples from 72 patients with a diagnosis of either Acute Myeloid Leukemia (AML) or Acute Lymphoblastic Leukemia (ALL). RNA was hybridised to Affymetrix microarrays containing 6817 human gene probes to generate the data. The dataset has served as a classic benchmarking dataset in many published computational methods (Li et al., 2011; Liao & Chin, 2007; Sweeney et al., 2015) and is available in processed and normalised form from https://www.kaggle.com/datasets/crawford/gene-expression.

Kaforou *et al*. (2013) collected blood samples of 584 adults in Malawi or South Africa with active Tuberculosis infection, latent Tuberculosis infection, or other disease. Transcriptional microarray profiles were generated using HumanHT-12 v.4 expression Beadarrays (Illumina). For FS-PLS benchmarking, we retained only the active TB (n = 195) or latent TB samples (n = 167) and used these for binary classification. Data is available for download from NCBI’s Gene Expression Omnibus under accession number GSE3725.

#### RNA-Seq datasets

The RNA-seq data from Ng *et al*. (2021) consists of 286 nasopharyngeal and 53 whole-blood samples from patients with COVID-19 and controls collected at University of California San Francisco. Sequencing of RNA samples was performed using the Illumina NovaSeq6000. For benchmarking, we used only nasopharyngeal samples and used the signature generation methods to classify COVID-19 (n = 138) from other acute viral respiratory illness (n = 120). RNA count tables were downloaded from the Gene Expression Omnibus under accession GSE163151.

The RAPIDS dataset was generated through the work of the Rapid Acute Paediatric Diagnosis of Infection in Suspected Sepsis study for the discovery of transcriptomic signatures for the diagnosis of bacterial vs viral sepsis in children (Schlapbach et al., 2024). Patients with suspected sepsis were recruited from Emergency Departments and Paediatric Intensive Care Units at four hospitals around Queensland, Australia. Whole blood samples were used for the study and split into a discovery cohort (n = 595) and a validation cohort (n = 316). RNA sequencing was performed on Illumina NovaSeq. Clinical diagnoses were confirmed and assigned to one of 6 classes: Definite Bacterial, Definite Viral, Probable Bacterial, Probable Viral, Non-infectious, or Unknown. For the benchmarking performed in this paper, Unknown cases were excluded and probable viral or bacterial cases were united with the definite diagnoses to construct a 3-class classification problem of Non-infectious vs Viral infection vs Bacterial infection.

Evaluation of models from RAPIDS data was performed using the validation cohort of the study.

#### Proteomic dataset

Álvez *et al*. (2023) produced a dataset of 1477 samples and 1463 protein features using the OLink Explore PEA technology. The samples originated from patients with 12 different cancer types and healthy controls, but for the FS-PLS benchmarking we selected the samples of the four blood cancer types (AML, CLL, DLBCL, Myeloma) and merged all other cancer samples into a control class (CTRL) to construct a 5-class classification task for assessment of the feature selection methods.

### Design of benchmarking experiment

The experiment is conducted as illustrated in Figure 1. Samples from each dataset are divided into 5 folds to allow cross-validation of model performance. Training samples are normalised independently from the test set and are used to select discriminative features for class prediction. For feature selection we compare FS-PLS with LASSO, Elastic Net, and MRMR as commonly used signature generation methods, and with ordinary forward selection methods (stagewise forward selection for binary datasets, stepAIC for multi-class datasets) to demonstrate FS-PLS’s improvement over its most related methods (see *Feature selection methods and settings*). The test set is then used to evaluate class performance. All methods use the same fold structure in the data to preserve comparability in results.

**Figure 1.**
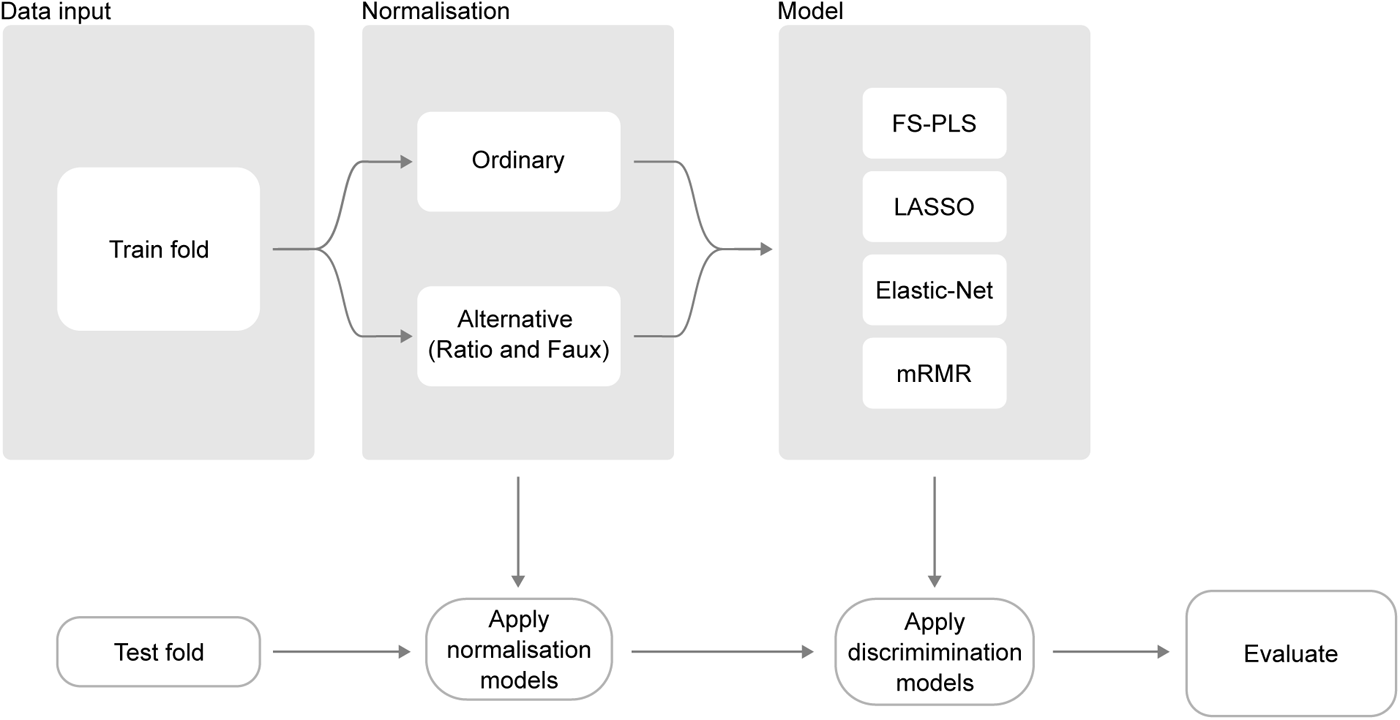
Schematic of the experimental setup for benchmarking of FS-PLS against LASSO, Elastic-Net, and mRMR. All datasets were normalised (ordinary normalisation) or, in the case of RNA-Seq data, modelled to reproduce log library size of samples with the fewest possible features and then normalised with the modelled solution (alternative normalisations). Following normalisation, we ran the feature discovery modelling to select features and learn a function to predict the outcome variable in question. Finally, models were evaluated by applying the various normalisations to test folds and generating new predictions using models trained on the respective normalisations. While Forward Stagewise and stepAIC methods were also tested in our benchmarking, there were not included in out alternative normalisation approach so are excluded from the schematic shown here.

Alongside the feature selection task, the training step for RNA-Seq data includes a *normalisation* feature discovery task where the log-library size of each sample is calculated and used as the continuous response variable for training of a sparse FS-PLS model to predict library size.

The computed normalisation model is fed into the discrimination feature discovery, where it contributes to two alternative normalisation solutions: “faux” normalisation based on predicted library size output by the normalisation model, and “ratio” normalisation where the original set of features is recast into a new feature space consisting of the log ratios of each feature and each normalising feature selected by the normalisation model.

The discrimination feature discovery task thus not only compares feature selection methods across folds, but also compares prediction accuracy from three possible normalisation approaches: full library size normalisation, faux normalisation, and ratio normalisation. Discrimination models are trained on the differently normalised data, yielding 12 total models to be evaluated.

For evaluation of performance, the held-out test partition is once again normalised with the three different normalisation solutions: full library size normalisation, faux normalisation by imputing a library size using the pre-trained model, and ratio normalisation using the earlier selected features as denominators in the feature log ratios. Trained discrimination models are then applied to generate new predictions and are assessed against true classes according to the appropriate metric (see *Methods*). Crucially, the fold structure used for normalisation and discrimination training is the same, so that new predictions are made on samples that have not been seen by either model in the training stage.

#### Data Treatment

For each of the 5 datasets in the benchmarking, samples were split into 5 folds for cross validation of model accuracy, thus yielding 5 individual models per method per dataset with 4 folds used for training and one fold held out for testing. To shorten computation time, pre-filtering of features was applied by selecting the top 10000 most variable features (Kaforou, Ng, RAPIDS) prior to splitting data into folds. For microarray data, where auto-scaling of features was used (mean = 0, variance = 1), scaling was performed separately within training and test partitions to prevent data leakage. RNA-Seq data was log transformed but left unscaled to ensure new predictions could be generated on individual samples. Missing values in the Álvez dataset were imputed within each fold using the *knnImpute* method from the *preProcess* function in the *caret* package.

The samples within each fold were identical for each of the methods tested to preserve comparability of results across methods. Furthermore, samples in the training fold for each model were kept consistent across feature selection for classification and for normalisation tasks, so that final classification predictions were determined on samples that had not been seen by either the classification or the normalisation model.

To construct sparse models for imputing library size, true library sizes of the test samples were determined by summing feature counts per sample before any filtering. The vectors of library sizes were log-transformed and used as a gaussian outcome variable for which the feature selection and model generation methods were modelled using gene counts *X*.

When incorporating feature selection for library size normalisation, it is important to ensure that the selected features are stably expressed in the discovery and the evaluation datasets. Accordingly, we added another filtering layer on features input into library size modelling so that only features in common across the discovery and test datasets’ 10000 most expressed features (.05 trimmed mean) were included. Gene counts were log-transformed but otherwise un-normalised and un-scaled before training. For the Ng data, the same 10000 most expressed used were used as candidates for normalisation and discrimination features. For the RAPIDS data, discrimination features were further selected from genes in common across the 10000 most variable features in the discovery and validation sets.

#### Feature selection methods and settings

We used the R packages *glmnet* (Friedman et al., 2010) and *mRMRe* (De Jay et al., 2013) for implementations of the alternative feature selection approaches. For LASSO and Elastic-Net (*glmnet)* an additional inner structure of 5 folds was used in the training data to select the strength of regularization (lambda). The implementation of MRMR (*mRMRe)* used in this study has no stopping criteria or model-fitting procedure and simply orders input features according to the MRMR loss function. Accordingly, we instructed MRMR to select the top *n* features equal to the number of features chosen by FS-PLS for that fold. We then estimated the coefficients of MRMR’s chosen features with ridge regression in *glmnet* and 5-fold cross-validation to select the degree of shrinkage.

The standard forward selection (FS) procedure holds some substantial similarities with FS-PLS. Stagewise FS proceeds with a series of iterations, at each point selecting a new feature to include in the model and adding new weights in addition to adjusting the weights of the features already selected (Efron et al., 2004). There are two primary differences between stagewise FS and FS-PLS. First, weights for selected features in FS-PLS are learned just once and then fixed in the deflated space with the aim of avoiding overfitting. Secondly, at each stage of the FS algorithm the current estimate is subtracted from the true class value to produce a residual outcome variable for selecting of the next feature, whereas FS-PLS employs the projection method of PLS to extract information already explained from the X matrix.

Given the similarities of the methods, it is prudent to evaluate FS-PLS in direct comparison with ordinary stagewise FS. We performed a head-to-head comparison of the two forward selection methods using the same 5-fold structure in each method for all datasets. For binary datasets, we used the *lars* package to generate stagewise FS models. However, stagewise FS is not applicable to multinomial data, as the current estimate at each stage cannot be subtracted from the outcome variable. To test an analogous method to stagewise FS in multinomial data, we used the *multinom* function from the *nnet* package in conjuction with *StepAIC* from the *MASS* package (Venables & Ripley, 2002) to build a sparse multinomial model to a pre-selected number of features.

Loss functions were selected for each type of response variable that was to be predicted: Area Under the Curve (AUC) for binary outcomes, multinomial deviance for multi-class outcomes, and Mean-Squared Error (MSE) for the continuous library sizes in the normalisation modelling. The same metrics were used in final performance evaluation, except for multi-class models whose performance is presented in multiclass AUC, and sensitivity and specificity per class.

For each method in multi-class and continuous prediction tasks, we extracted two sets of coefficients: one which produced the minimum training error (‘min’), and another which gave the maximum regularization strength that still produced training error within 1 standard error of the minimum (‘1se’). In *glmnet*, this is performed automatically with the *cv.glmnet* function, but in FS-PLS we achieved the equivalent by directly selecting the number of features that gave the min and 1se training errors. We found that the set of features corresponding to the minimum training error generally performed best, so all reported classification results are from ‘min’ models apart from the normalisation feature selection models for which the ‘1se’ models are used.

#### Analysis environment and code availability

Analysis was performed in R version 4.2.1. Visualisation of results was achieved using *ggplot2* version 3.4.4, with assistance from *ROCR* (Sing et al., 2005) version 1.0.11 and *pROC* (Robin et al., 2011) version 1.18.4 for deriving metrics, and *caret* (Kuhn, 2008) version 6.0.94 for partition splitting and missing value imputation in proteomic data.

FS-PLS is available for the community to use at https://github.com/lachlancoin/fspls. Scripts for creation and evaluation of results discussed in this paper are deposited at https://github.com/dn-ra/FSPLS-publication-repo.

## Results

### FS-PLS generates much smaller signatures compared with regularisation methods

We first assess FS-PLS in the setting in which the library size (or equivalent) information is available, for example when the test set is measured on the same platform as the training dataset.

We observed that FS-PLS generally achieved similar performance to LASSO and Elastic Net but with much fewer features selected for inclusion in the model (Figure 2). In two binary datasets, there was no significant loss in performance from LASSO to FS-PLS according to mean difference in AUC (Kaforou: +0.01, Ng: -0.008) despite a many-fold reduction in mean feature numbers (Ng: 4.4 (FS-PLS), 31.4 (LASSO); Kaforou: 6 (FS-PLS), 19.6 (LASSO)).

**Figure 2.**
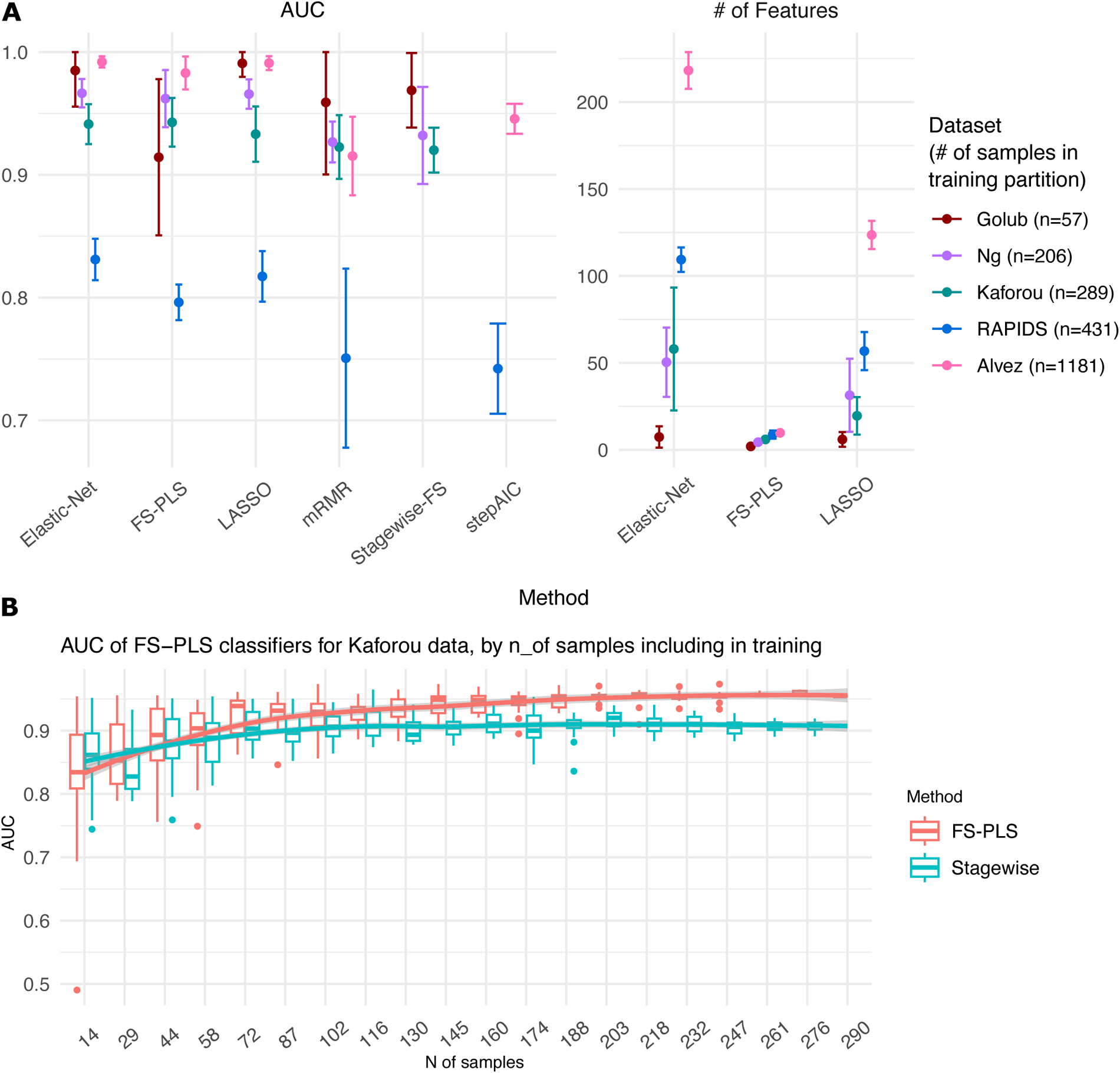
Performance of examined methods on each dataset. **A.** Test AUCs (left) and numbers of features selected (right) for all datasets and methods included in the study. For most datasets, FS-PLS performs similarly to LASSO and Elastic-Net with comparatively much fewer features. Mean and 95% confidence interval across each of 5-folds are shown. For multiclass datasets (Rapids, Alvez), AUC value for each k-fold is derived from the average AUC across classes. Display of feature numbers selected excludes mRMR, Stagewise-FS, and stepAIC as these methods were instructed to select the same number of features as chosen by FS-PLS. Only one of Stagewise-FS or stepAIC are applied to each dataset, as they are the most appropriate forward selection strategies for binary or multinomial problems respectively. **B.** Performance of FS-PLS against ordinary Stagewise forward selection on Kaforou data at varying sample sizes to explain poor FS-PLS results in Golub dataset. Each box represents 20 repeated samplings of the data at the given sample size. Stagewise FS initially displays greater accuracy than FS-PLS at low sample sizes but is surpassed by it as more samples are included in the training set.

The Golub dataset diverged from the above results by demonstrating a sharp drop in FS-PLS performance in contrast with the other methods (Figure 2A). The results seen here are best explained by the small number of samples available in the dataset and will likely be recoverable by training on a larger dataset. We demonstrate this below in our comparison of FS-PLS with ordinary forward selection methods.

Moreover, as the sample size of the datasets increased, the reduction of feature numbers became more pronounced. Application of regularisation-based methods to multi-class problems led to several hundred features selected. In contrast, FS-PLS created a much smaller signature (mean # of features: 10 (RAPIDS dataset), 7.6 (Álvez dataset)) at a cost to multiclass AUC of 5-6 percentage points (Figure 2, Supplementary tables).

### FS-PLS outperforms MRMR for the equivalent number of selected features

Since MRMR was set to select the same number of features as FS-PLS, this represents a biased advantage for MRMR as it is borrowing the appropriate number of features learned by FS-PLS. Despite this, MRMR and FS-PLS selected different features to include in the model’s set and delivered varying performance (Figure 2). Aside from the exception of the Golub dataset, as discussed above, FS-PLS outperforms MRMR binary classification models when trained and tested on common folds (+0.02, +0.03 mean AUC for Kaforou and Ng, respectively).

In multi-class datasets, FS-PLS outperforms again but is also more sensitive to under-represented classes than MRMR (Figure 3). When weighting samples by their inverse class proportions for the RAPIDS dataset, performance of FS-PLS on all 3 classes actually deteriorates compared with no sample weights at all. Conversely, MRMR requires sample weights for 2 out of 3 classes to achieve best performance and is outperformed by FS-PLS in the aggregate.

**Figure 3.**
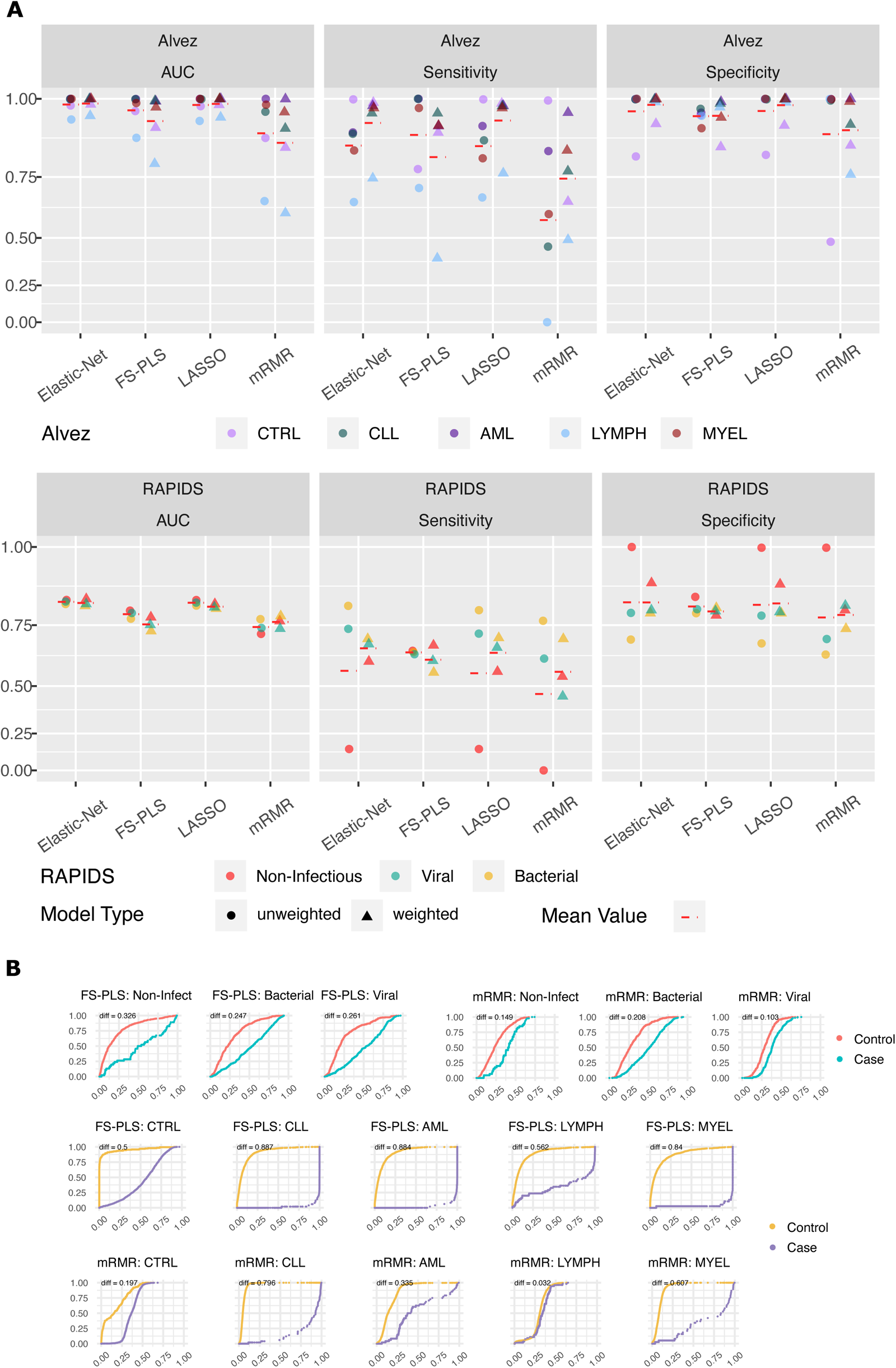
Model accuracy from FS-PLS and competing methods on multiclass datasets where each class is assessed separately. Predictions from test folds within each method are concatenated to provide an overall measure of prediction accuracy for each class. For both RAPIDS (A) and Alvez (B) datasets, model accuracy from both unweighted (left) and weighted models (right) are shown, and performance is presented in AUC, Sensivity, and Specificity per class. For Alvez data, unweighted FS-PLS models were trained through to a maximum number of 10 features regardless of training performance because heavy class imbalance led to deterioration of training loss with the inclusion of any more than one variable. Overall, prediction accuracy from FS-PLS is better in unweighted than weighted models, and vice versa for the competing methods. Hence, our continued discussion of FS-PLS’s multiclass performance refers to unweighted FS-PLS models in comparison with the weighted versions derived from LASSO, Elastic-Net and mRMR. B. Cumulative probability curves for sample prediction of each class in the two multi-class datasets from FS-PLS’s unweighted and mRMR’s weighted model. FS-PLS models maintain much greater separation between classes than mRMR as indicated by the area between coloured lines. Lines are coloured by case or control where cases are samples with true class corresponding to the named label of the column and controls are samples from the other classes. Curves are annotated with Wasserstein distance between controls and cases, where higher values indicate more confidence separation of the two by the model.

Training FS-PLS without sample weights also performed poorly for the Álvez data. The control class is so dominant in number that the inclusion of any more than one variable leads to a deterioration of the training loss and the number of features selected never exceeds one. To bypass the model training difficulty, we trained an FS-PLS model for each fold through to a maximum number of 10 features regardless of training performance. As seen with the RAPIDS dataset, the unweighted FS-PLS Álvez models demonstrate superior or equal test AUC results for every class compared with the weighted MRMR models (Figure 3). The benefit of sample weighting for mRMR models is unclear in the AUC results when compared with unweighted models. However, the meagre sensitivity of the LYMPH class (0.0) without weights is justification of adopting the weighted model as superior. Indeed, all alternative models suffer from meagre sensitivity for one class in both multinomial datasets when sample weights are unused.

Given FS-PLS’s sensitivity to low proportion classes in the absence of weighting, it is unclear whether a more optimal strategy than the one used here may further improve performance. Nevertheless, the results of feature selection on multi-class datasets reported throughout this benchmarking take the unweighted FS-PLS models as compared with weighted models from the alternative methods.

In addition to comparing methods using AUC, sensitivity, and specificity, we implemented a Wasserstein metric to quantify the separation in cumulative predicted probabilities for each binary comparison in the multinomial classification problem (see *Methods)*. Visualising the divergence of probabilities provides an assessment of prediction performance that accounts for the clearance each method gives to candidate classes that is not visible in AUC calculations.

The difference in case vs control cumulative probability curves for FS-PLS is much higher than for MRMR, indicating a relatively greater degree of confidence in the probabilities being assigned (Figure 3). Indeed, the unweighted FS-PLS model on the RAPIDS data exhibits a greater separation in probabilities for each class than for all other weighted methods (Supplementary Figure 1). Altogether, these results indicate that FS-PLS is a method that delivers high performance discrimination models with relatively few features chosen.

### FS-PLS generally outperforms standard stagewise forward selection methods on the tested datasets

As discussed previously, FS-PLS exhibits poor performance on the relatively simple Golub binary dataset, and so stagewise FS has superior accuracy on test partitions by an average of +0.055 AUC (Figure 2). In other binary datasets, however, FS-PLS outperforms stagewise FS using the same number of features (mean AUC change Ng +0.03, Kaforou +0.02).

We hypothesised that the poor performance of FS-PLS models from the Golub data are a consequence of the small number of samples used in training these (n = 57 per model). We trialled the theory by performing a sampling experiment on the Kaforou microarray dataset. The data was split into an 80%-20% train-test partition. Random sampling of the observations in the train partition was performed at increments of 5% between 5% and 100% of the total partition size. Sampling was repeated 20 times within each sample size. FS and FS-PLS models were trained on the sampled datasets, ultimately yielding 400 models (20 sample sizes, 20 replicates within each) across the spectrum of sample sizes that were then evaluated on the held-out test partition.

The modelling results (Figure 2) demonstrate that at low sample numbers, models built with ordinary FS perform better than FS-PLS. With increasing sample size, FS-PLS accuracy improves more rapidly than ordinary FS and ultimately outperforms it at mid-high sample sizes. With such limited sample numbers in the Golub dataset, the poor performance of FS-PLS is likely due to the performance differential demonstrated in this sampling experiment.

For multinomial datasets, FS-PLS achieves higher AUC than stepAIC in all classes for both RAPIDS and Álvez data (Figure 2). The improved performance of FS-PLS over stepAIC in multinomial datasets make it a promising method for this use as there are few methods available for sparse multinomial feature selection. Furthermore, it is simpler to construct than StepAIC, as the latter is burdened by the need to construct a full linear model using all features before proceeding through feature selection steps.

### FS-PLS selects features for normalisation that retain accuracy without requiring a true normalisation factor

Next we addressed the problem of library size normalisation. The development of sparse signatures for diagnosis from omics data is generally hampered by the requirement to measure all the molecules in the assay for normalisation of the data (eg. library size). We demonstrate a novel use of FS-PLS to learn normalising features in the RNA-Seq datasets which scale reliably and strictly with the dataset’s library size. By using these features for normalisation, we bypass the need to survey all molecules in the assay and raise the possibility of generating diagnosis signatures that are constituted entirely of a set of discriminating features plus a small selection of features for normalisation of the data.

Using the Ng and RAPIDS datasets, we program FS-PLS to progressively choose features and fit coefficients to reproduce the log library size of each sample until the log-likelihood model of the model is no longer significant against the null model (see *Methods).* A 5-fold structure was used again so that five models were produced for each dataset and performance could be evaluated on the respective hold-out group.

For most datasets and folds, the algorithm halted after two features had been selected and so for simplicity we limited the number of normalising features to two globally. The adjusted R^2^ value for the correlation between true and predicted log-library sizes in the held-out folds was 0.92 for Ng and 0.88 for RAPIDS (Supplementary Figure 2), indicating a high predictive accuracy. We used the selected normalising features in two ways: firstly, to normalise feature counts in each sample by the predicted log-library size, which we call ‘Faux’ normalisation; and secondly, to generate a new feature matrix constituted of each feature vector divided by the feature vector of each normalisation feature, which we call ‘Ratio’ normalisation. With 2 normalisation features in a fold, the Ratio normalisation approach leads to a new data matrix of dimension *n* x *2p*. Our implementation of a ratio normalisation was inspired by the work of Wang *et al*. (2022) in their Cross-Platform Omics Prediction (CPOP) procedure.

We employed both the alternative normalisation approaches in a new discrimination feature selection round and compared the classification performance of the resulting models against models computed on the ordinarily normalised data. Selection of discriminating features was re-performed with all methods and the same train-test partitions were used for normalisation feature selection to ensure that samples in the held-out folds had never before been encountered by either the normalisation or discrimination model generation.

Figure 4 shows the results of the experiment with alternative normalisation strategies. The Ng data once again delivers similar performance from LASSO, Elastic-Net, FS-PLS and mRMR across the various normalisation strategies. The similar performance is a promising finding, given that the ratio and faux normalisation solutions from FS-PLS now only require an average of 3.6 discrimination features for faux normalisation and 5.8 discrimination features for ratio normalisation, plus two additional normalisation features, to classify a new sample. The result, if replicated in wider cohorts, could present an opportunity for conversion of the omics signature into a point-of-care diagnostics test.

**Figure 4.**
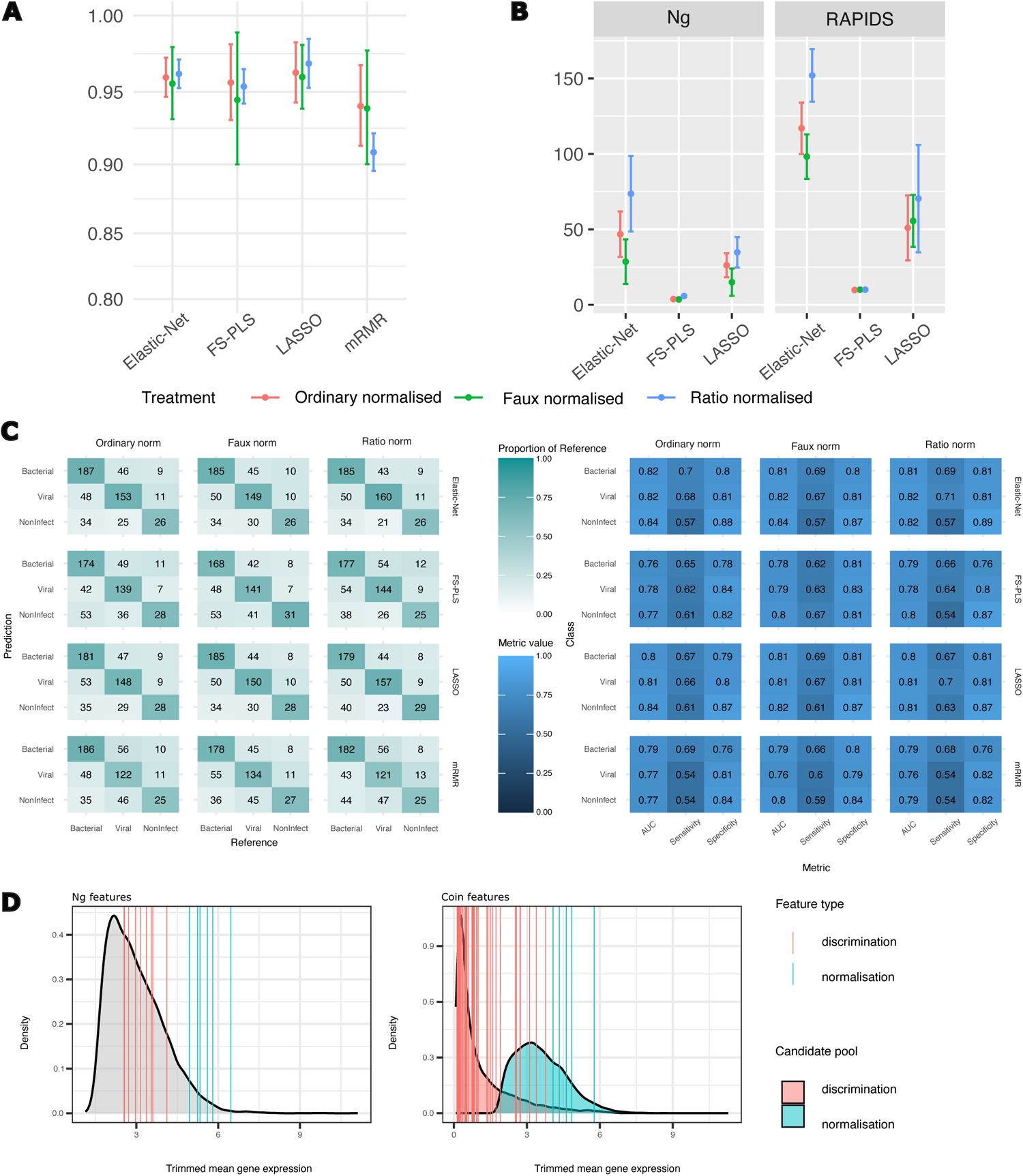
Results of sparse normalisation approaches on RNA-Seq datasets. **A.** Mean and 95%CI AUC results for each fold of Ng data when models constructed using ordinary or alternative normalisation strategies. Normalisation features were selected with FS-PLS before being used to build class discrimination models with LASSO, Elastic Net, FS-PLS, and mRMR. Broadly, FS-PLS’s alternative normalisations produce comparable accuracy results to the full ‘ordinary’ normalisation. **B.** Numbers of features selected for the models displayed in A and C. Count excludes the features used in normalisation (n=2 for Faux and Ratio normalisation). As in Figure 2A, number of mRMR features is excluded as it is explicitly instructed to match FS-PLS. What is exchanged for the accuracy differences in A and C is a complete model with few features needed for normalisation and discrimination of samples. **C.** Concatenated results for class predictions from all 5 folds of RAPIDS data when models constructed on varying normalisation strategies. Left tile plot shows confusion matrix of predictions per method and per normalisation approach with predicted class on the vertical axis. Right plot shows AUC, sensitivity and specificity per class. As above, the compressed ‘faux’- and ‘ratio’-normalised modelling approaches display only slim accuracy differences while discarding the need for full library size calculation. D. Discrimination and normalisation features from FS-PLS Faux normalisation models (5 folds per dataset) displayed according to their trimmed mean expression in the original datasets and superimposed over density plots of the trimmed mean for all features considered. Selected normalisation features tend to be expressed at much higher levels than discrimination features. In the RAPIDS dataset, normalisation and discrimination features were selected from different subsets of the features (10000 highest trimmed mean expressed, and 10000 most variably expressed genes, respectively), so density plots are split according to the different pools. The two feature types were chosen from the same feature subset in the Ng data (10000 highest trimmed mean expressed).

In the RAPIDS dataset, Elastic-Net and LASSO generally outperform the other methods in each normalisation approach, as might be expected given the results that have already been revealed in this study. mRMR and FS-PLS show little difference in accuracy. Once again, however, the great reduction in number of features needed for the model makes the use of FS-PLS an attractive prospect for development of a diagnostic test. The total signature size from FS-PLS is 10 discrimination features plus 2 normalisation features on each fold, compared with an average of 50 (plus 2 normalisation features) for LASSO. The 5-fold reduction in feature numbers comes at a small AUC cost of 0.02 in both alternative normalisation models (0.81 multiclass AUC LASSO, 0.79 multiclass AUC FS-PLS).

### Performance of alternatively normalised models on validation data mirrors performance of ordinary-normalised dataset

To further evaluate the use of FS-PLS for coupled normalisation and classification, we trained FS-PLS on the entire RAPIDS dataset and tested its performance on an external validation set (Figure 7). As with the cross-validation setup described above, we used FS-PLS to select features predictive of sample log-library size, then proceeded with training FS-PLS to predict diagnosis class from ordinary, faux, or ratio normalised RNA-seq data. All models selected a set of 12 features: 10 for discrimination and 2 for normalisation.

Prediction results on the validation data suffered a severe drop in fidelity (Figure 5), primarily due to a reduced performance on new Non-Infectious samples (0.9 train AUC vs 0.7 validation AUC; Ordinary normalisation). Recovery of the Non-Infectious class has been a difficult task in earlier studies on sepsis biomarkers (Pierrakos & Vincent, 2010) and, indeed, the ratio-normalised model mis-classified every Non-Infectious sample as Bacterial or Viral. Yet the decline in performance from Training to Validation data is consistent across the Ordinary and Faux normalised models, suggesting that the elimination of the full library normalisation does not inject any further error into the model. Furthermore, models trained with LASSO and Elastic-

**Figure 5.**
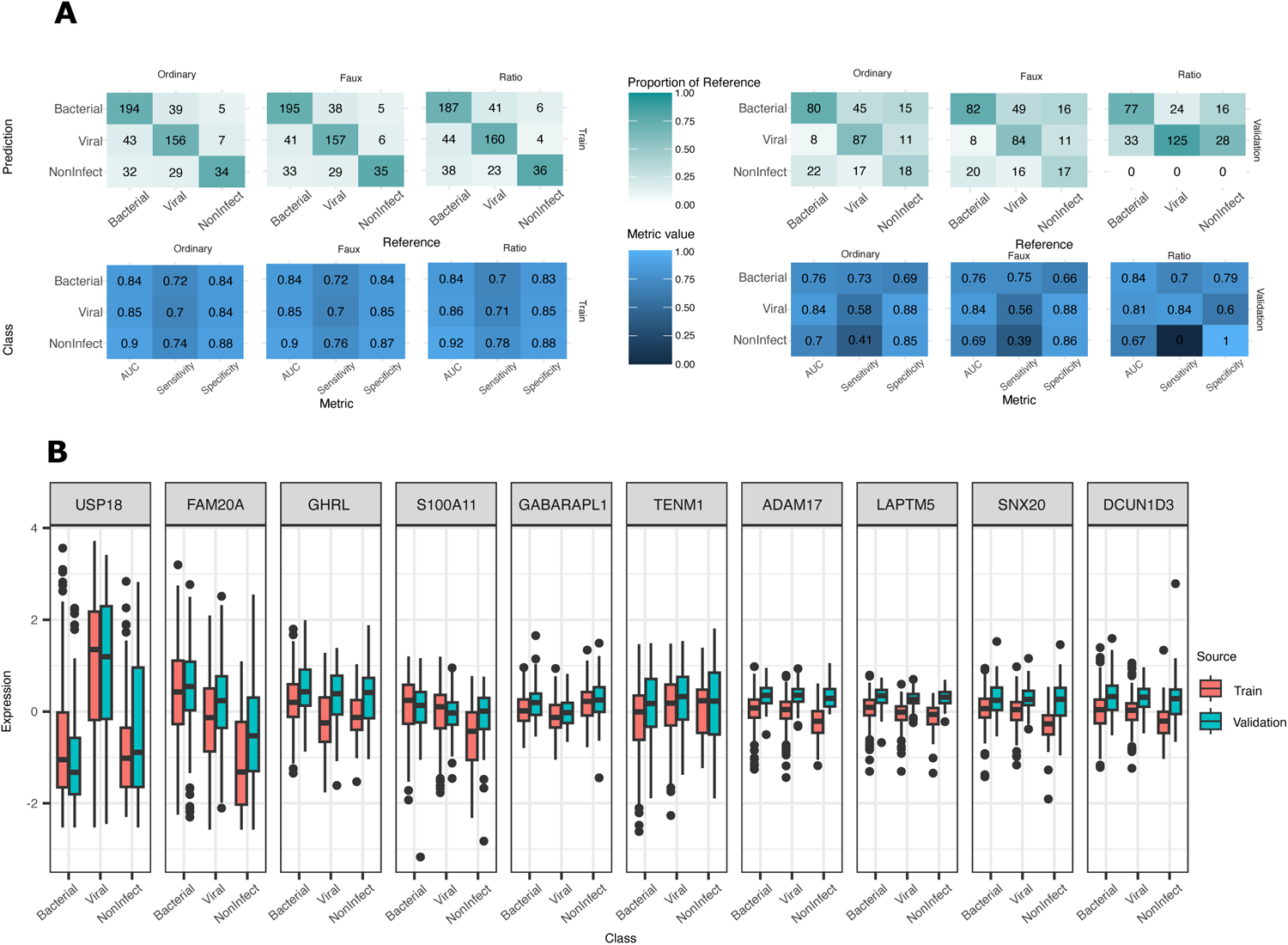
Performance assessment of FS-PLS on RAPIDS validation data. **A.** Performance of FS-PLS models on RAPIDS train and validation data. The left side displays training error, right side displays validation error. The top row of plots are confusion matrices, with each square coloured by the proportion of reference class captured by the prediction. The bottom row of plots displays AUC, Sensitivity, and Specificity metrics for each class. Results are additionally separated according to the normalisation approach used. Model accuracy suffers a sizeable drop in accuracy when applied to validation data and is consistent across normalisation types. **B.** Expression profiles of selected features across training and validation dataset, separated into true class of each sample. Data is from ‘ordinary’ normalisation approach and shows unmatched distributions across the datasets for many of the chosen features. The drop in validation performance shown in A may be related to the shifted distribution of normalised expression values since validation data is centred according to mean feature values learned in the training data.

Net on the three alternatively normalised datasets also show comparable validation performance with FS-PLS (Elastic-Net Ordinary-normalisation validation AUCs: 0.82 (Bacterial), 0.83 (Viral), 0.69 (Non-Infectious), Supplementary Figure 3, Supplementary tables), and the loss of validation accuracy is more likely attributable to the broader issue of generalisability across experimental batches (Figure 5, see *Discussion*). Since the expanded feature set used in these models might be expected to deliver close to maximum accuracy, the close performance from FS-PLS is a positive result that demonstrates the feasibility of the truly sparse FS-PLS model.

## Discussion

In this work we have benchmarked and formally described an effective method for biomarker discovery, FS-PLS, which derives small predictive signatures of disease. We have demonstrated the flexibility and applicability of the method using five publicly available datasets across three ‘omics technologies. In all datasets, FS-PLS produces signatures with a small number of features and high predictive performance in comparison with widely used alternatives.

FS-PLS will be useful for researchers wishing to derive predictive signatures of disease that require as few features as possible. We anticipate the accelerated development of point-of-care lab-on-a-chip (LOC) diagnostic devices in the coming decade, for which minimal signatures of disease are required. In such a scenario, FS-PLS is well placed to deliver features for conversion into an LOC. We have demonstrated that signatures generated with FS-PLS underperform those from LASSO and Elastic-Net in cross-fold validation. However, the trade-off of size to performance is one that will prompt researchers to use FS-PLS over alternatives in applications that require a small signature.

In this study we also demonstrate the usefulness of FS-PLS for selecting features which act to normalise the measurement of the discrimination features. It remains unknown whether normalisation features detected from a high-throughput assay will also prove effective in a condensed LOC test. It may be that such features reflect not a biological control for variation but rather a technical artefact of the assay, in which case their suitability in an alternative technology will be muted. Alternatively, Oxford Nanopore Technologies’ targeted sequencing shows potential for miniaturisation and diagnostic specialisation (Kang et al., 2020; World Health Organisation, 2023). With a technology such as this that has consistent mode of operation in both full and reduced form, determining a normalisation feature using a full assay may only require re-evaluation of the normalisation weights for it to be applied in the specialised application. However, existing demonstrations of rapid targeted sequencing tests have searched for sequence variations (SNPs, structural variants) or binary presence/absence of a genetic feature rather than exploring quantitative gene signatures (Tabata et al., 2022; Zhang et al., 2023).

There are several possible ways the work presented here may be improved. Firstly, the generalisability of FS-PLS models to new data needs to be demonstrated further. There is difficulty in obtaining separate discovery and validation datasets that follow identical sample and library processing steps. When validating our RAPIDS dataset, we aimed to test the model on an earlier dataset (Habgood-Coote et al., 2023), but found that variations in the protocol led to poor agreement of the feature expression in the two sets. Likewise, an attempted test of our normalisation strategy using a library model built on an external dataset (Lieberman et al., 2020) failed to reproduce an accurate library size in the Ng data because of a ribosomal RNA depletion step in the library preparation. A more robust demonstration of FS-PLS’s end-to-end signature generation should use a larger number of datasets with more aligned discovery and validation protocols.

Aside from protocol differences, it is likely that ordinary batch effects are also contributing to the validation performance drop off seen in the RAPIDS data. Our approach intends to create fully online-capable signatures – processing samples individually without any information gleaned from a wider sample set. As such, each feature is centralised by the mean of the feature learned from the training data, which for many features fails to truly centralise the validation data (Figure 5C). Training FS-PLS on a larger set of data from more sources will help to minimise the influence of dataset-specific patterns and produce more generalisable signatures. However, the question of how to reliably scale individual samples is an open one without a clear solution.

We have identified several computational improvements that could be made to the algorithm. Firstly, the program would benefit from a more efficient method of selecting the next most important feature. Least Angle Regression (LARS) (Efron et al., 2004) is one such method, and in development tests was effective at speeding up analysis of binary datasets. However, there is no trivial way to extend LARS to multinomial data, so in this setting we are at present limited to the relatively slow procedure of fitting a multinomial linear model for every variable at every selection step. Earlier research has demonstrated some potential for the implementation of LARS in a multinomial scenario by using a non-canonical link function (Gluhovsky, 2012). Future research on FS-PLS should investigate ways by which LARS might be extended or approximated for multinomial outcomes.

Modellers of biological data often seek to predict samples on an ordinal scale like disease severity (e.g. mild, moderate, severe) or progression (e.g. early, middle, late). For these outcomes it is clear there is an order to the categories. To make FS-PLS useful for these types of data, existing procedures for feature selection in ordinal data should be investigated for adaptation into our method (Baccianella et al., 2010; Mukras et al., 2007). On this, the advantage in the design of FS-PLS when incorporating new methods is that no adjustments are made to the *y* vector at each feature inclusion. So long as there are means by which to select individually relevant features, the process can be repeated to include more features after each successive deflation of the *X* matrix. Indeed, the deflation of *X* gives FS-PLS a significant advantage over ordinary FS methods for multinomial problems as it avoids the requirement to subtract the estimate ỹ at each iteration which cannot be performed for categorical data. For this reason, feature selection on multinomial data is achieved with alternative methodologies like *stepAIC* which requires modelling of the full feature space and, as we show above, is outperformed by FS-PLS.

Finally, FS-PLS would benefit from alternative options for stopping criteria. At present, the algorithm stops when the log likelihood of additional features ceases to achieve significant difference from the null model. The criteria permit a new feature to enter the model that is statistically significant even when the additional accuracy it delivers is low. Alternatively, instituting a cross-fold error stopping criteria would allow the algorithm to halt its search when the added feature no longer improves performance in held out samples, thus leading to signatures that increase in size only if they benefit accuracy.

## Supporting information

Supplementary Figures and Extended Pseudocode

Supplementary Results Tables

## Acknowledgements

This research was supported by The University of Melbourne’s Research Computing Services and the Petascale Campus Initiative. D.R. was supported by the Australian Government Research Training Programme (RTP) scholarship. L.C. was supported by the National Health and Medical Research Council Grant GNT1195743.

## References

Altman, N., & Krzywinski, M. (2018). The curse(s) of dimensionality. Nature Methods, 15(6), 399–400. 10.1038/s41592-018-0019-x

Álvez, M. B., Edfors, F., von Feilitzen, K., Zwahlen, M., Mardinoglu, A., Edqvist, P. H., Sjöblom, T., Lundin, E., Rameika, N., Enblad, G., Lindman, H., Höglund, M., Hesselager, G., Stålberg, K., Enblad, M., Simonson, O. E., Häggman, M., Axelsson, T., Åberg, M., . . . Uhlén, M. (2023). Next generation pan-cancer blood proteome profiling using proximity extension assay. Nat Commun, 14(1), 4308. 10.1038/s41467-023-39765-y

Baccianella, S., Esuli, A., & Sebastiani, F. (2010). *Feature selection for ordinal regression* Proceedings of the 2010 ACM Symposium on Applied Computing, Sierre, Switzerland. 10.1145/1774088.1774461

Beaude, A., Rafiee Vahid, M., Augé, F., Zehraoui, F., & Hanczar, B. (2023). AttOmics: attention-based architecture for diagnosis and prognosis from omics data. Bioinformatics, 39(Supplement_1), i94–i102. 10.1093/bioinformatics/btad232

Cai, J., Luo, J., Wang, S., & Yang, S. (2018). Feature selection in machine learning: A new perspective. Neurocomputing, 300, 70–79. 10.1016/j.neucom.2017.11.077

Cui, H., Wang, C., Maan, H., Pang, K., Luo, F., Duan, N., & Wang, B. (2024). scGPT: toward building a foundation model for single-cell multi-omics using generative AI. Nature Methods. 10.1038/s41592-024-02201-0

De Jay, N., Papillon-Cavanagh, S., Olsen, C., El-Hachem, N., Bontempi, G., & Haibe-Kains, B. (2013). mRMRe: an R package for parallelized mRMR ensemble feature selection. Bioinformatics, 29(18), 2365–2368. 10.1093/bioinformatics/btt383

Ding, C., & Peng, H. (2005). Minimum redundancy feature selection from microarray gene expression data. J Bioinform Comput Biol, 3(2), 185–205. 10.1142/s0219720005001004

Efron, B., Hastie, T., Johnstone, I., & Tibshirani, R. (2004). Least angle regression. The Annals of Statistics, 32(2), 407–499. 10.1214/009053604000000067

Friedman, J. H., Hastie, T., & Tibshirani, R. (2010). Regularization Paths for Generalized Linear Models via Coordinate Descent. Journal of Statistical Software, 33(1), 1–22. 10.18637/jss.v033.i01

Gliddon, H. D., Herberg, J. A., Levin, M., & Kaforou, M. (2018). Genome-wide host RNA signatures of infectious diseases: discovery and clinical translation. Immunology, 153(2), 171–178. 10.1111/imm.12841

Gliddon, H. D., Kaforou, M., Alikian, M., Habgood-Coote, D., Zhou, C., Oni, T., Anderson, S. T., Brent, A. J., Crampin, A. C., Eley, B., Heyderman, R., Kern, F., Langford, P. R., Ottenhoff, T. H. M., Hibberd, M. L., French, N., Wright, V. J., Dockrell, H. M., Coin, L. J., . . . Levin, M. (2021). Identification of Reduced Host Transcriptomic Signatures for Tuberculosis Disease and Digital PCR-Based Validation and Quantification. Front Immunol, 12, 637164. 10.3389/fimmu.2021.637164

Gluhovsky, I. (2012). Multinomial least angle regression. IEEE Trans Neural Netw Learn Syst, 23(1), 169–174. 10.1109/tnnls.2011.2178480

Golub, T. R., Slonim, D. K., Tamayo, P., Huard, C., Gaasenbeek, M., Mesirov, J. P., Coller, H., Loh, M. L., Downing, J. R., Caligiuri, M. A., Bloomfield, C. D., & Lander, E. S. (1999). Molecular classification of cancer: class discovery and class prediction by gene expression monitoring. Science, 286(5439), 531–537. 10.1126/science.286.5439.531

Habgood-Coote, D., Wilson, C., Shimizu, C., Barendregt, A. M., Philipsen, R., Galassini, R., Calle, I. R., Workman, L., Agyeman, P. K. A., Ferwerda, G., Anderson, S. T., van den Berg, J. M., Emonts, M., Carrol, E. D., Fink, C. G., de Groot, R., Hibberd, M. L., Kanegaye, J., Nicol, M. P., . . . Kaforou, M. (2023). Diagnosis of childhood febrile illness using a multi-class blood RNA molecular signature. Med, 4(9), 635–654.e635. 10.1016/j.medj.2023.06.007

Herberg, J. A., Kaforou, M., Wright, V. J., Shailes, H., Eleftherohorinou, H., Hoggart, C. J., Cebey-Lopez, M., Carter, M. J., Janes, V. A., Gormley, S., Shimizu, C., Tremoulet, A. H., Barendregt, A. M., Salas, A., Kanegaye, J., Pollard, A. J., Faust, S. N., Patel, S., Kuijpers, T., . . . Consortium, I. (2016). Diagnostic Test Accuracy of a 2-Transcript Host RNA Signature for Discriminating Bacterial vs Viral Infection in Febrile Children. Jama, 316(8), 835–845. 10.1001/jama.2016.11236

Jackson, H. R., Zandstra, J., Menikou, S., Hamilton, M. S., McArdle, A. J., Fischer, R., Thorne, A. M., Huang, H., Tanck, M. W., Jansen, M. H., De, T., Agyeman, P. K. A., Von Both, U., Carrol, E. D., Emonts, M., Eleftheriou, I., Van der Flier, M., Fink, C., Gloerich, J., . . . Zwerenz, M. (2023). A multi-platform approach to identify a blood-based host protein signature for distinguishing between bacterial and viral infections in febrile children (PERFORM): a multi-cohort machine learning study. The Lancet Digital Health, 5(11), e774–e785. 10.1016/S2589-7500(23)00149-8

Kaforou, M., Wright, V., Oni, T., French, N., & Anderson, S. (2013). Detection of Tuberculosis in HIV-Infected and-Uninfected African Adults Using Whole.

Kang, A. S. W., Bernasconi, J. G., Jack, W., & Kanavarioti, A. (2020). Ready-to-use nanopore platform for the detection of any DNA/RNA oligo at attomole range using an Osmium tagged complementary probe. Scientific Reports, 10(1), 19790. 10.1038/s41598-020-76667-1

Kuhn, M. (2008). Building Predictive Models in R Using the caret Package. Journal of Statistical Software, 28(5), 1–26. 10.18637/jss.v028.i05

Lê Cao, K.-A., Boitard, S., & Besse, P. (2011). Sparse PLS discriminant analysis: biologically relevant feature selection and graphical displays for multiclass problems. BMC Bioinformatics, 12(1), 253. 10.1186/1471-2105-12-253

Li, S., Harner, E. J., & Adjeroh, D. A. (2011). Random KNN feature selection - a fast and stable alternative to Random Forests. BMC Bioinformatics, 12, 450. 10.1186/1471-2105-12-450

Li, Y., Mansmann, U., Du, S., & Hornung, R. (2022). Benchmark study of feature selection strategies for multi-omics data. BMC Bioinformatics, 23(1), 412. 10.1186/s12859-022-04962-x

Liao, J. G., & Chin, K.-V. (2007). Logistic regression for disease classification using microarray data: model selection in a large p and small n case. Bioinformatics, 23(15), 1945–1951. 10.1093/bioinformatics/btm287

Lieberman, N. A. P., Peddu, V., Xie, H., Shrestha, L., Huang, M. L., Mears, M. C., Cajimat, M. N., Bente, D. A., Shi, P. Y., Bovier, F., Roychoudhury, P., Jerome, K. R., Moscona, A., Porotto, M., & Greninger, A. L. (2020). In vivo antiviral host transcriptional response to SARS-CoV-2 by viral load, sex, and age. PLoS Biol, 18(9), e3000849. 10.1371/journal.pbio.3000849

Mabey, D., Peeling, R. W., Ustianowski, A., & Perkins, M. D. (2004). Diagnostics for the developing world. Nature Reviews Microbiology, 2(3), 231–240. 10.1038/nrmicro841

Meinshausen, N. (2007). Relaxed Lasso. Computational Statistics & Data Analysis, 52(1), 374–393. 10.1016/j.csda.2006.12.019

Mukras, R., Wiratunga, N., Lothian, R., Chakraborti, S., & Harper, D. (2007). Information gain feature selection for ordinal text classification using probability re-distribution. Proceedings of the Textlink workshop at IJCAI,

Ng, D. L., Granados, A. C., Santos, Y. A., Servellita, V., Goldgof, G. M., Meydan, C., Sotomayor-Gonzalez, A., Levine, A. G., Balcerek, J., Han, L. M., Akagi, N., Truong, K., Neumann, N. M., Nguyen, D. N., Bapat, S. P., Cheng, J., Martin, C. S., Federman, S., Foox, J., . . . Chiu, C. Y. (2021). A diagnostic host response biosignature for COVID-19 from RNA profiling of nasal swabs and blood. Sci Adv, 7(6). 10.1126/sciadv.abe5984

Park, T., & Casella, G. (2008). The Bayesian Lasso. Journal of the American Statistical Association, 103(482), 681–686. 10.1198/016214508000000337

Pierrakos, C., & Vincent, J.-L. (2010). Sepsis biomarkers: a review. Critical Care, 14(1), R15. 10.1186/cc8872

Robin, X., Turck, N., Hainard, A., Tiberti, N., Lisacek, F., Sanchez, J. C., & Müller, M. (2011). pROC: an open-source package for R and S+ to analyze and compare ROC curves. BMC Bioinformatics, 12, 77. 10.1186/1471-2105-12-77

Saeys, Y., Inza, I., & Larrañaga, P. (2007). A review of feature selection techniques in bioinformatics. Bioinformatics, 23(19), 2507–2517. 10.1093/bioinformatics/btm344

Schlapbach, L. J., Ganesamoorthy, D., Wilson, C., Raman, S., George, S., Snelling, P. J., Phillips, N., Irwin, A., Sharp, N., Le Marsney, R., Chavan, A., Hempenstall, A., Bialasiewicz, S., MacDonald, A. D., Grimwood, K., Cling, J. C., McPherson, S. J., Blumenthal, A., Kaforou, M., . . . Coin, L. (2024). Host gene expression signatures to identify infection type and organ dysfunction in children evaluated for sepsis: a multicentre cohort study. The Lancet Child & Adolescent Health. 10.1016/S2352-4642(24)00017-8

Sing, T., Sander, O., Beerenwinkel, N., & Lengauer, T. (2005). ROCR: visualizing classifier performance in R. Bioinformatics, 21(20), 3940–3941. 10.1093/bioinformatics/bti623

Sweeney, T. E., Chen, A. C., & Gevaert, O. (2015). Combined Mapping of Multiple clUsteriNg ALgorithms (COMMUNAL): A Robust Method for Selection of Cluster Number, K. Scientific Reports, 5(1), 16971. 10.1038/srep16971

Tabata, Y., Matsuo, Y., Fujii, Y., Ohta, A., & Hirota, K. (2022). Rapid detection of single nucleotide polymorphisms using the MinION nanopore sequencer: a feasibility study for perioperative precision medicine. JA Clinical Reports, 8(1), 17. 10.1186/s40981-022-00506-7

Tibshirani, R. (1996). Regression Shrinkage and Selection via the Lasso. Journal of the Royal Statistical Society. Series B (Methodological), 58(1), 267–288. http://www.jstor.org/stable/2346178

Venables, B., & Ripley, B. (2002). Modern Applied Statistics With S. In. 10.1007/b97626

Wang, K. Y. X., Pupo, G. M., Tembe, V., Patrick, E., Strbenac, D., Schramm, S.-J., Thompson, J. F., Scolyer, R. A., Muller, S., Tarr, G., Mann, G. J., & Yang, J. Y. H. (2022). Cross-Platform Omics Prediction procedure: a statistical machine learning framework for wider implementation of precision medicine. npj Digital Medicine, 5(1), 85. 10.1038/s41746-022-00618-5

World Health Organisation. (2023). *Use of targeted next-generation sequencing to detect drug-resistant tuberculosis* [Rapid communication]. https://iris.who.int/bitstream/handle/10665/371687/9789240076372-eng.pdf?sequence=1

Xu, C., & Jackson, S. A. (2019). Machine learning and complex biological data. Genome Biology, 20(1), 76. 10.1186/s13059-019-1689-0

Zhang, Y., Lu, X., Tang, L. V., Xia, L., & Hu, Y. (2023). Nanopore-Targeted Sequencing Improves the Diagnosis and Treatment of Patients with Serious Infections. MBio, 14(1), e0305522. 10.1128/mbio.03055-22

Zou, H. (2006). The Adaptive Lasso and Its Oracle Properties. Journal of the American Statistical Association, 101(476), 1418–1429. http://www.jstor.org/stable/27639762

Zou, H., & Hastie, T. (2005). Regularization and variable selection via the elastic net. Journal of the Royal Statistical Society: Series B (Statistical Methodology), 67(2), 301–320. 10.1111/j.1467-9868.2005.00503.x

