## Supplementary Figures and Extended Pseudocode for "A flexible framework for minimal biomarker signature discovery from clinical omics studies without library size normalisation"

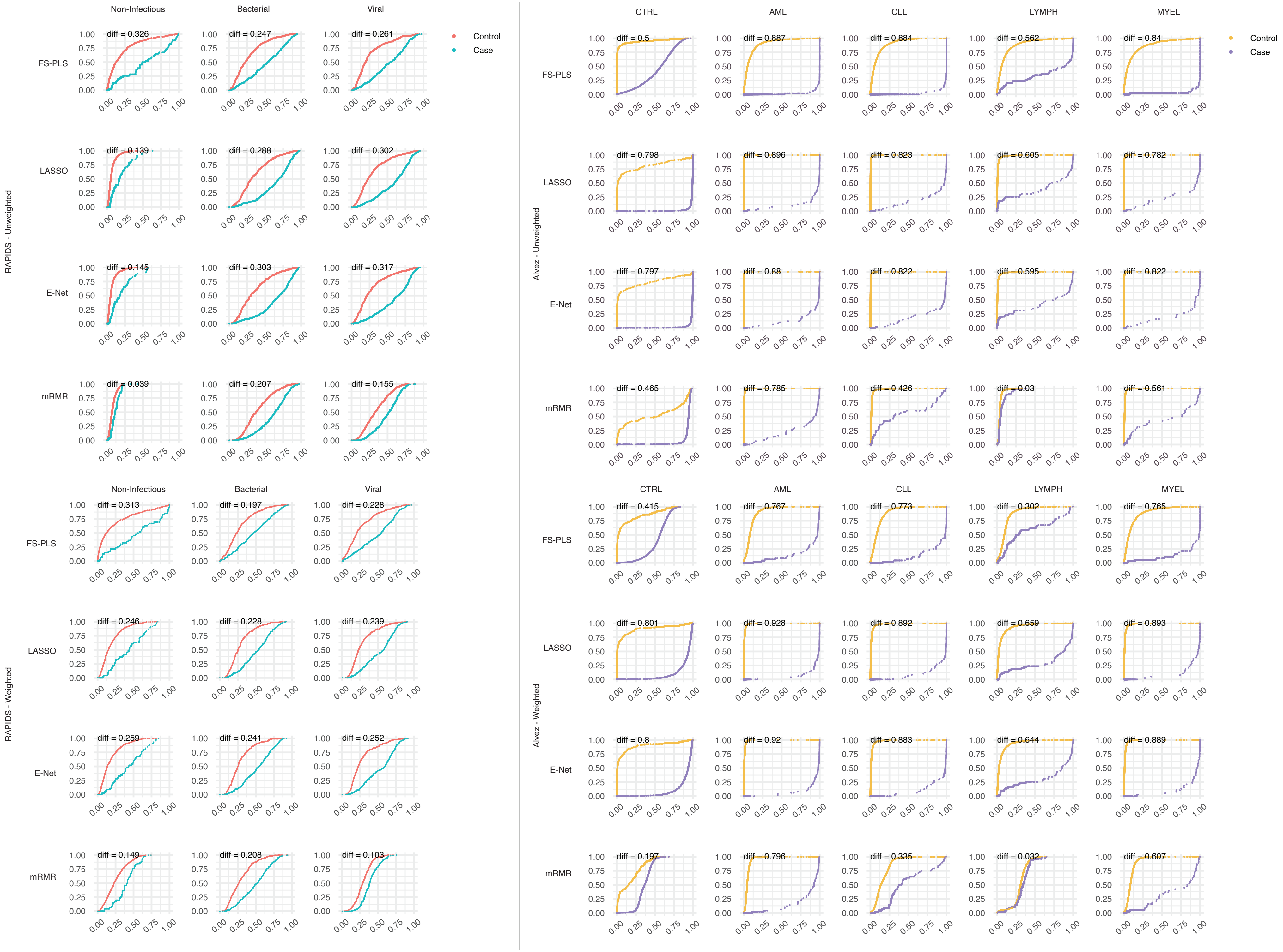
RAPIDS - Unweighted
Non-Infectious
Bacterial
Viral
Control
Case
FS-PLS
diff = 0.326
diff = 0.247
diff = 0.261
LASSO
diff = 0.139
diff = 0.288
diff = 0.302
E-Net
diff = 0.145
diff = 0.303
diff = 0.317
mRMR
diff = 0.039
diff = 0.207
diff = 0.155
Alvez - Unweighted
CTRL
AML
CLL
LYMPH
MYEL
Control
Case
FS-PLS
diff = 0.5
diff = 0.887
diff = 0.884
diff = 0.562
diff = 0.84
LASSO
diff = 0.798
diff = 0.896
diff = 0.823
diff = 0.605
diff = 0.782
E-Net
diff = 0.797
diff = 0.88
diff = 0.822
diff = 0.595
diff = 0.822
mRMR
diff = 0.465
diff = 0.785
diff = 0.426
diff = 0.03
diff = 0.561
RAPIDS - Weighted
Non-Infectious
Bacterial
Viral
FS-PLS
diff = 0.313
diff = 0.197
diff = 0.228
LASSO
diff = 0.246
diff = 0.228
diff = 0.239
E-Net
diff = 0.259
diff = 0.241
diff = 0.252
mRMR
diff = 0.149
diff = 0.208
diff = 0.103
Alvez - Weighted
CTRL
AML
CLL
LYMPH
MYEL
FS-PLS
diff = 0.415
diff = 0.767
diff = 0.773
diff = 0.302
diff = 0.765
LASSO
diff = 0.801
diff = 0.928
diff = 0.892
diff = 0.659
diff = 0.893
E-Net
diff = 0.8
diff = 0.92
diff = 0.883
diff = 0.644
diff = 0.889
mRMR
diff = 0.197
diff = 0.796
diff = 0.335
diff = 0.032
diff = 0.607

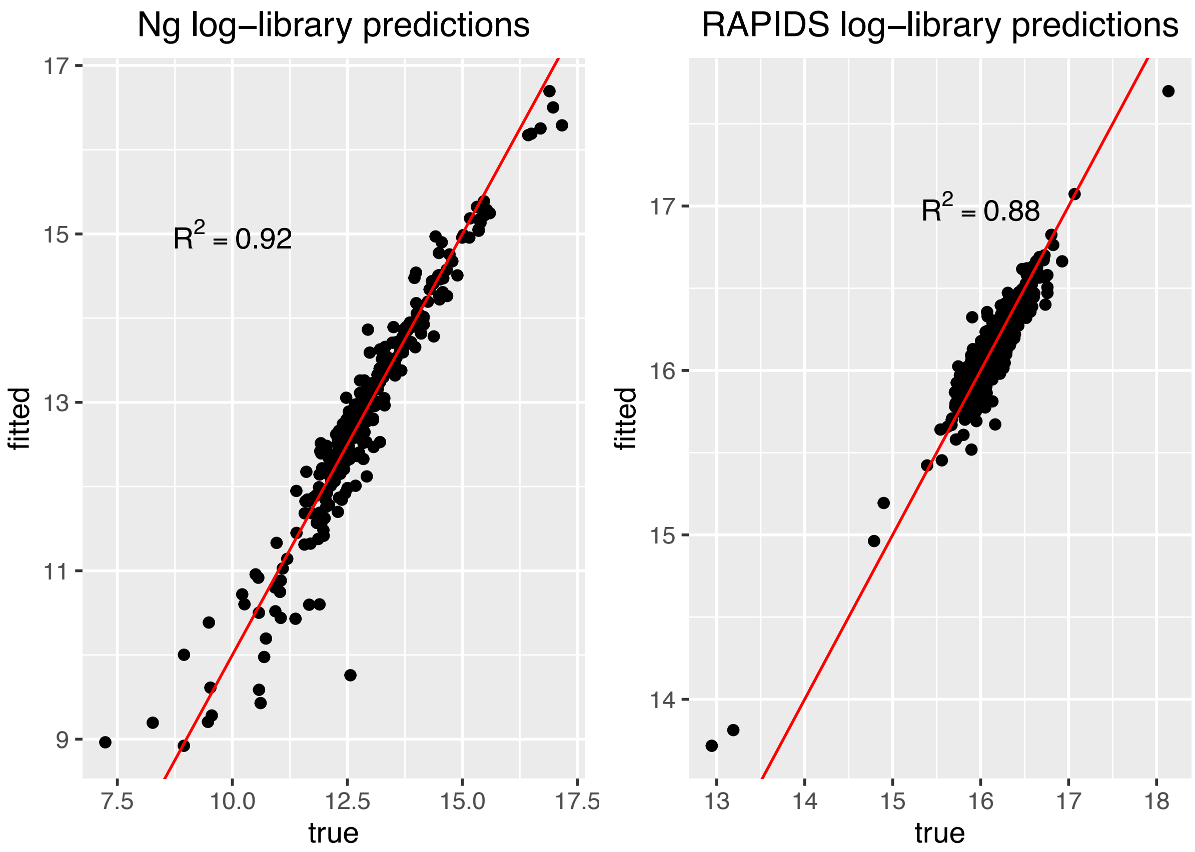


Supplementary Figure 2.

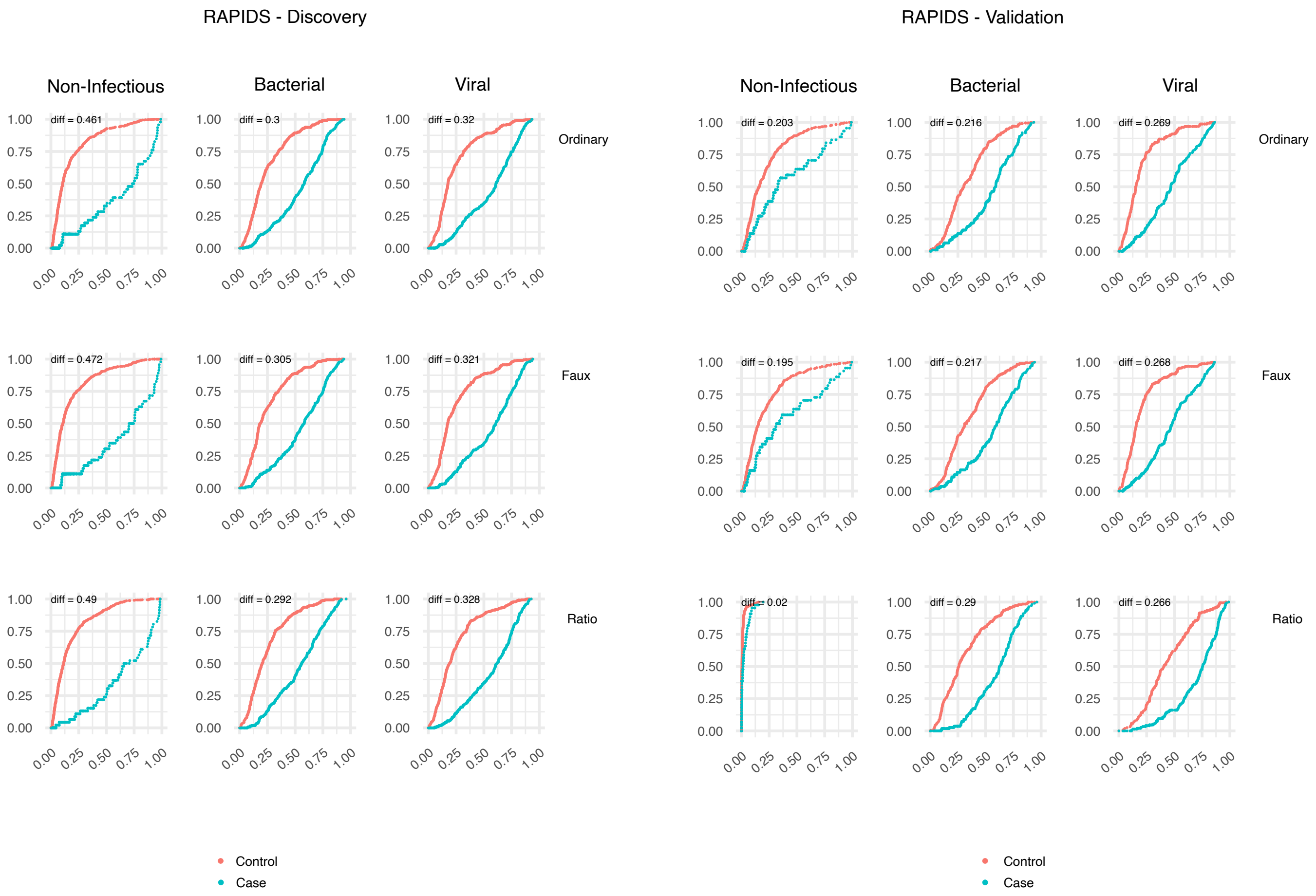

RAPIDS - Discovery
RAPIDS - Validation
Non-Infectious
Bacterial
Viral
Ordinary
Faux
Ratio
diff = 0.461
diff = 0.3
diff = 0.32
diff = 0.472
diff = 0.305
diff = 0.321
diff = 0.49
diff = 0.292
diff = 0.328
diff = 0.203
diff = 0.216
diff = 0.269
diff = 0.195
diff = 0.217
diff = 0.268
diff = 0.02
diff = 0.29
diff = 0.266
0.00
0.25
0.50
0.75
1.00
Control
Case

### Pseudocode of FS-PLS algorithm:

```
FUNCTION FeatureSelection(matrix X, vector y, threshold T, maxVariables M) ->
variables, betas, transformations
    INIT matrix Xk with 0 columns and X.rows rows
    INIT integer k = 0
    INIT betas
    INIT transformations W

    // Calculate and subtract column means from X
    FOR j from 1 to X.columns
        mean = average of the j-th column of X
        Subtract mean from each element in the j-th column of X
    EndFOR

    WHILE k < M
        // Singular Value Decomposition on Xk
        U, D, V.transpose() = SVD(Xk)

        // Project X onto Xk
        Pk = U * U.transpose() * X

        // Deflate X with Pk
        Rk = X - Pk

        INIT maxLogLikelihood = -infinity
        INIT bestJ = -1

        // Fit a univariate linear model for each column of Rk
        FOR j from 1 to Rk.columns
            model = fitLinearModel(Rk[:, j], y)
            logLikelihood = calculateLogLikelihood(model)

            IF logLikelihood > maxLogLikelihood
                maxLogLikelihood = logLikelihood
                bestJ = j
            EndIF
        EndFOR

        // Fit coefficient with L2 shrinkage (ridge regression)
        coefficient = fitWithRidgeRegression(Rk[:, bestJ], y)

        // Add the selected variable to Xk
        Xk = concatenate(Xk, Rk[:, bestJ] multiplied by coefficient)

        // Add coefficients to betas list
        betas = concatenate(betas, coefficient)

        // Add transformation to W list
        proj = U.tranpose() * X
        new_W = V.inverse() * D.inverse() * proj
        W = concatenate(W, new_W)


        // Chi-squared test
        pValue = chiSquaredTest(logLikelihood(model_with_Xk),
                                logLikelihood(null_model))
```

```
        IF pValue > T
            BREAK
        EndIF

        k = k + 1
    EndWHILE

    SET variables = column_names(Xk)

    RETURN variables, betas, W
EndFUNCTION
```

**Supplementary Figure 1.**
Cumulative probability curves for sample prediction of unweighted (top) vs weighted (bottom) models for each method on multi-class datasets RAPIDS (left) and Alvez (right). Greater separation of lines, measured by Wasserstein distance in the top left of each plot, indicates greater confidence in the predictions made by each model for the named class. Lines are coloured by case or control where cases are samples with true class corresponding to the named label of the column and controls are samples from the other classes. Weighting of samples by inverse proportion during training generally adds confidence in prediction for LASSO and Elastic-Net, but FS-PLS models perform best when no sample weighting is used.

**Supplementary Figure 2.**
Log-library size predictions (y-axis) against true values (x-axis) for Ng and RAPIDS datasets. Red line marks the diagonal where x=y. A cross-validation approach was used so that new predictions are made by models that have not seen the same data during training.

**Supplementary Figure 3.**
Cumulative probability curves, as in Supplementary Figure 1, for FS-PLS models on discovery (left) and validation (right) cohorts of RAPIDS data. Rows separate normalisation solutions used in the models: ordinary (top), faux (middle), and ratio (bottom). All normalisation approaches experience a drop in accuracy when applied to validation data. The ratio-normalised model loses the ability to identify non-infectious sample, but the faux-normalised model suffers no further drop in performance than ordinary normalisation.
